# Contrasting responses of control and fibrotic lung fibroblasts to fibrotic stimuli: the role of osteoprotegerin in extracellular matrix remodeling

**DOI:** 10.1101/2025.02.20.639137

**Authors:** Yanzhe Liu, Mehmet Nizamoglu, Fenghua Zhao, Theo Borghuis, Taco Koster, Prashant K. Sharma, Martin C. Harmsen, Matthew J. Thomas, Eric S. White, Karim C. El Kasmi, Irene H Heijink, Barbro N. Melgert, Janette K. Burgess

## Abstract

Idiopathic pulmonary fibrosis (IPF) is a progressive and fatal lung disease characterized by dysregulated wound healing, leading to excessive production of extracellular matrix (ECM) proteins, particularly collagen, by activated fibroblasts. Osteoprotegerin (OPG), a soluble protein associated with ECM remodeling in bone, has been found in higher levels in serum and lung tissue of IPF patients compared to controls. However, whether OPG is associated with ECM remodeling by fibroblasts in lung fibrosis remains poorly understood. In this study we assessed differences in OPG levels in ECM-derived hydrogels from non-disease controls and patients with IPF encapsulated with primary human lung fibroblasts from control and patients with IPF (7 days). OPG levels were assessed in a collected culture medium and stained lung tissue sections, and related to biomechanical properties.

Fibrotic human lung fibroblasts secreted and deposited more OPG compared to controls, which was observed in both non-diseased and IPF ECM hydrogels. OPG levels correlated significantly with matrix stiffness and collagen fiber structural properties, but only in IPF hydrogels, independent of fibroblast source. Strikingly, we also observed opposite correlations for control and fibrotic fibroblasts in fibrotic hydrogels. Control fibroblasts within a fibrotic microenvironment, secreted less OPG when hydrogel stiffness increased and deposited more OPG when collagen fibers were more aligned. Interestingly, fibrotic fibroblasts secreted more OPG when collagen fiber length increased and when collagen fiber curvature increased. Overall, our findings suggest that control and fibrotic lung fibroblasts exhibited opposite behavior with respect to OPG and ECM remodeling under fibrotic conditions and this may have consequences for matrix stiffness

## Introduction

A normal repair process in tissue involves differentiation of fibroblasts into myofibroblasts and their subsequent activation. These myofibroblasts are major producers of extracellular matrix (ECM)^1^. Consequently, ECM proteins, such as collagens, are deposited in and around the injury site to restore the architecture of tissue. Once the architecture has been reestablished, ECM production is downregulated and excess scar tissue is remodeled^2^. In idiopathic pulmonary fibrosis (IPF) the wound-healing process is dysregulated. IPF is a progressive and fatal lung disease characterized by fibroblast foci in lung tissue with excessively activated fibroblasts within the interstitial space^3^. These activated fibroblasts continue to produce ECM proteins unchecked, leading to progressive stiffening and dysfunction of lung tissue and ultimately an irreversible scarring (fibrosis) in the lung^4^.

A key hallmark of lung fibrosis is the progressive rigidity of the lung, driven by enhanced ECM crosslinking and increased mechanical resistance (stiffness). This high stiffness provides active mechanical cues driving behavior of cells within the tissue^5,6,7^. In addition to matrix stiffness, altered collagen fiber structure is also a dominant driver of cellular actions in disease conditions. Aligned collagen can drive cells to elongate earlier and faster than collagen with random alignment^8^ and it also enhances cell motility^9^ in tumor microenvironments. Moreover, aligned collagen can lead to greater matrix stiffness compared to unaligned collagen^10^, suggesting that ECM stiffness and altered collagen structure can drive cell behavior. A recent study in our group^11^ has found that non-fibrotic and fibrotic lung fibroblasts exhibited active and different behaviors in response to stimuli of a fibrotic microenvironment, particularly with respect to ECM stiffness and fiber structure. Another study found that control dermal fibroblasts were capable of remodeling a gradient ECM-derived hydrogel by decreasing stiffness in the high stiffness zone^12^. These findings highlight the bidirectional interactions between fibroblasts and fibrotic ECM, characterized by changes in collagen fiber organization and ECM stiffness. In turn, fibroblasts exhibit various adaptive responses to a fibrotic environment by remodeling ECM properties. Osteoprotegerin (OPG) is a soluble protein that plays an essential role in regulating ECM for maintaining bone integrity^13^. Recently, we showed that more OPG is present in serum and lung tissue from patients with IPF compared to those non-diseased controls, and it could be a potential biomarker of early fibrotic activity in lung and liver^14,15,16^. Notably, OPG is not only a secreted protein with a recognized role as a decoy receptor for receptor activator of nuclear factor kappa-B ligand (RANKL), but can also be anchored into ECM via binding to various matrix proteins. These OPG binding partners include connective tissue growth factor (CTGF)^17^, fibronectin, and basement membrane ECM components such as collagen IV as well as laminin^18^. However, little is known about an association between OPG and mechanical properties of ECM including stiffness and fiber structure in lung fibrosis.

In this study we hypothesized that the fibrotic lung microenvironment stimulates lung fibroblasts to produce high levels of OPG, increasing matrix stiffness and creating aligned fibers in a fibrotic context. To address this hypothesis, we established a three-dimensional (3D) in vitro lung fibrosis model using control and IPF decellularized lung ECM derived hydrogels and encapsulated these with control and IPF lung fibroblasts. We then assessed levels of soluble and deposited OPG in this model. Based on our group^’^s previous finding that lung fibroblasts reorganize ECM as a response to fibrotic stimuli, we further investigated whether OPG produced by lung fibroblasts correlated with ECM structure and stiffness of ECM within non-fibrotic and fibrotic microenvironments.

## Methods and Materials

### Human lung fibroblasts encapsulated in dECM-derived hydrogels

The samples from lung decellularized ECM (dECM)-derived hydrogels in this study were generated in a previous study, reported in Nizamoglu et al^11^. The characteristics describing these hydrogel properties and ECM fiber properties were originally reported in this previous study and have been reanalyzed herein to relate them to the OPG levels measured in this study. Briefly, lung tissues from controls without lung fibrosis and patients with IPF were decellularized using a mixed decellularization protocol^5^. Next, these decellularized lung samples were lyophilized before being ground to fine lung ECM powders. Pepsin (2 mg/mL in 0.01 M HCI) digestion was applied to 40 mg dry ECM power using in a 7.5 mL glass vial to obtain solubilized lung ECM, followed by pH adjustment with 1M NaOH and the addition of 1 tenth volume of 10X phosphate-buffered saline (PBS) (Gibco, Grand Island, USA) solution to bring the solution back to physiological salts and ions to form a pre-gel. Primary human lung fibroblasts from either control or patients with IPF were mixed with the pre-gel solution, before it was allowed to gelate for 1-2 hours at 37C°. Cell growth media was added to these fibroblast encapsulated lung-ECM derived hydrogels and the samples were maintained for 7 days in an incubator (5 % CO2, 37°C), with a half growth media change on day 4. Supernatants were collected at media changes and at the experiment end for further testing.

### Enzyme-linked immune sorbent assay (ELISA)

Secreted OPG in supernatant was measured using ELISA (cat#DY805, R&D Systems Minneapolis, MN, USA) according to the instructions provided by the manufacturer. Samples were diluted 1:10 in reagent diluent.

### Immunohistochemistry

The hydrogels were fixed with 2% paraformaldehyde before being embedded in agarose (Invitrogen, Waltham, MA, USA) to prevent dehydration and embedded in paraffin. Five μm paraffin-embedded sections from human lung-ECM derived hydrogels were prepared and used for histological analysis. Sections were deparaffinized and rehydrated and antigen retrieval was performed with 10 mM citrate buffer at pH 6 at 100°C for 15 min. Endogenous peroxidases were blocked by incubation with 0.3% hydrogen peroxidase (H2O2) in PBS at room temperature for 30 min. After washing with 1X PBS three times, unspecific targets in sections were blocked by incubation with 4% (bovine serum albumin) BSA at room temperature for 30 min. Subsequently, OPG antibody (ab183910, Abcam, UK, 1:300 in lung ECM-derived hydrogel sections, was incubated on the sections overnight at 4°C. After washing with PBS, slides were incubated with secondary antibody goat anti-rabbit immunoglobulin-HRP in lung ECM-derived hydrogel (1:100, P0488 Dako, Denmark), for 45 minutes at RT. Slides were then washed three times in PBS, followed by three washes in demineralized water. Then Vector®NovaRED® (Vector Laboratories, California, United States) Substrate was added to the sections for 10 min to visualize the specific protein. After washing sections in demi water three times, they were counter stained with hematoxylin for 2 mins. Then the sections were mounted with histological mounting media (SP15100, Thermo Fisher Scientific, United States) and the slides were scanned using 40x objective on the Hamamatsu NanoZoomer digital slides scanner (Hamamatsu Photonic K.K).

### Imaging analyses

The digital images were opened and extracted as TIFF file using Aperio Imagescope (v12.3.3.5048, Leica Biosystems, Amsterdam, Netherlands). Adobe Photoshop 2024 (San Jose, California, USA) were performed to remove staining artifacts. ImageJ-win64 (LOCI, University of Wisconsin)^19^ was used to split images into red and blue channels respectively to represent the images stained with OPG (Novared image) and stained with hematoxylin and thresholds selected using appropriate vectors as previously determined^20^. All thresholds were used in a macro calculating tissue area per channel. The strength of the signal from each pixel was determined and categorized as “weak”, “moderate”, and “strong” based on level of the strength. After running all images, signals of pixels were categorized based on levels of intensity values in R studio (Boston, MA, USA) and sorted for the percentage of pixels of positive stained area per image. The calculation of the percentage positively stained area was performed using equation as shown below.

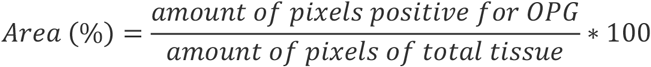

The measurement of the OPG intensity of the cells within the hydrogel was performed in Cellprofiler. Firstly, the staining images were split in ImageJ-win64 into three channels, including the image with Novared stain, hematoxylin stain and an image containing background. These images were inverted in Cellprofiler and used to measure the OPG positivity of the cells within the hydrogel. To enhance the cell features in the inverted grayscale and NovaRed image the enhance operation and feature type Texture was used within the Cellprofiler module EnhanceOrSupressFeatures. The resulting images were then stretched to the full intensity range with the module RescaleIntensity. Artefacts were then removed from the rescaled grayscaled image by subtracting it with the objects detected in the inverted background image. The background objects were detected using the IdentifyPrimaryObjects module with objects diameter set between 15 and 40 pixels. Before subtracting the detected background, objects were expanded by 9 pixels to ensure all background was removed. Next objects were detected in the background corrected grayscales image and the NovaRed rescaled image using the IdentifyPrimaryObjects module with objects diameter set between 15 and 60 pixels and objects diameter set between 25 and 75 pixels respectively. Next the objects found within the NovaRed image were masked by the objects found in the grayscale image. After shrinking these objects by 10 pixels, these objects were used input objects for the module identyfySecundaryObjects using the inverted NovaRed image. These objects were denoted as NovaRed positive cells. The intensity and area of these objects were measured in the NovaRed inverted image with the modules.

### Mechanical and structural measurements of hydrogels

The measurements of parameters of collagen fibers and stiffness were described previously^11^. Briefly, Stiffness was measured using Low Load Compression Tester^21^. Young’s Modulus was used to describe the value of stiffness. In this study, we calculated the change in stiffness in the hydrogel from day 0 to day 7. We used the stiffness value in the hydrogel encapsulated with cells on day 7 to subtract the stiffness value in hydrogels without cells. Parameters of collagen fibers were analyzed in lung tissue sections stained with picrosirius red using The Workflow Of Matrix BioLogy Informatics (TWOMBLI) performed in ImageJ^22^.

### Data and Statistical analyses

All statistical analyses were performed using GraphPad Prism v10.2.1 (GraphPad Company, San Diego, CA, US). QQ plots were used to explore the normality. Two-way analysis of variance (ANOVA) was used to analyze levels of secreted and deposited OPG, and also to check the effects of fibroblasts and lung ECM hydrogel separately and to identify when these two factors were interacting. A two-way ANOVA examines whether fibrotic lung ECM hydrogel changes OPG production compared to control lung ECM hydrogel, whether fibrotic lung fibroblasts change OPG production compared to control lung fibroblasts, and whether the effects of ECM hydrogel origin and fibroblast origin have a significant interaction. In case of a significant interaction, the results for separate factors cannot be trusted and a Šidák-corrected post hoc test was used to compare individual groups to explain the interaction. Parametric data from two-way ANOVA are shown as mean, p< 0.05 is considered statistically significant. Correlations between secreted and deposited OPG and ECM fiber characteristics, and the correlation between OPG intensity and changes of ECM stiffness in lung dECM hydrogels in which lung fibroblasts were encapsulated were assessed using a Pearson test for parametric data or a Spearman test for nonparametric data.

## Results

### More OPG secretion by fibroblasts derived from patients with IPF

To identify the effect of a fibrotic microenvironment on fibroblasts with respect to OPG production, we first measured secreted OPG in supernatants collected from control and IPF-derived lung fibroblasts encapsulated in control and IPF-derived lung-ECM hydrogels (Figure 1). Our results indicate that fibroblasts derived from donors with IPF secreted significantly more OPG compared to fibroblasts from control donors, regardless of whether they were encapsulated in control or IPF hydrogels.

**Figure 1:**
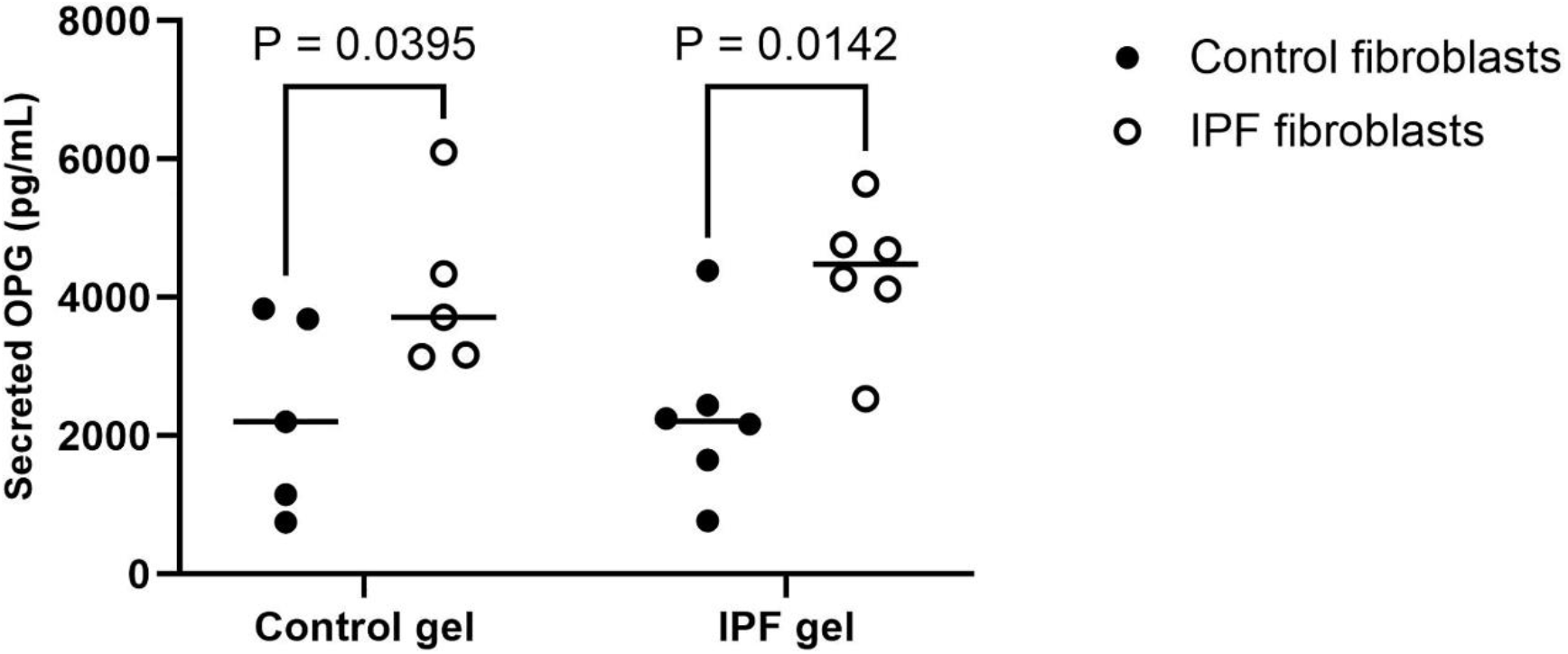
Secreted OPG in culture medium of control and IPF fibroblasts encapsulated in lung dECM-derived hydrogels. Control (n=5) and IPF (n=6) lung fibroblasts were encapsulated in control or IPF lung dECM-derived hydrogels for 7 days. Culture medium was collected to measure secreted OPG levels. Data analysed using two-way ANOVA with Šidák-corrected post hoc test, p< 0.05 was considered statistically significant. OPG: osteoprotegerin, IPF: idiopathic pulmonary fibrosis, dECM: decellularized extracellular matrix.

### More OPG deposition by fibroblasts derived from patients with IPF

Subsequently, we performed immunohistochemical staining of OPG in sections from our hydrogels to explore the effect of a fibrotic microenvironment on fibroblasts with respect to OPG deposition (Figure 2). We observed OPG located to both control and fibrotic lung fibroblasts either in control or IPF lung dECM hydrogels (Figure 2A), but we did not observe specific staining for OPG in empty hydrogels (data not shown). We calculated total intensity of OPG per cell (Figure 2B) and the percentage area of each hydrogel that was occupied by OPG (Figure 2C). Consistent with the pattern for secreted OPG, deposited OPG showed the same trend for fibrotic fibroblasts, regardless of the hydrogel type in which these fibroblasts were embedded, this was not seen to the same extent. This finding indicates that lung fibroblasts from patients with IPF are intrinsically different from controls and have high deposited OPG, which is not modulated by the surrounding matrix microenvironment.

**Figure 2:**
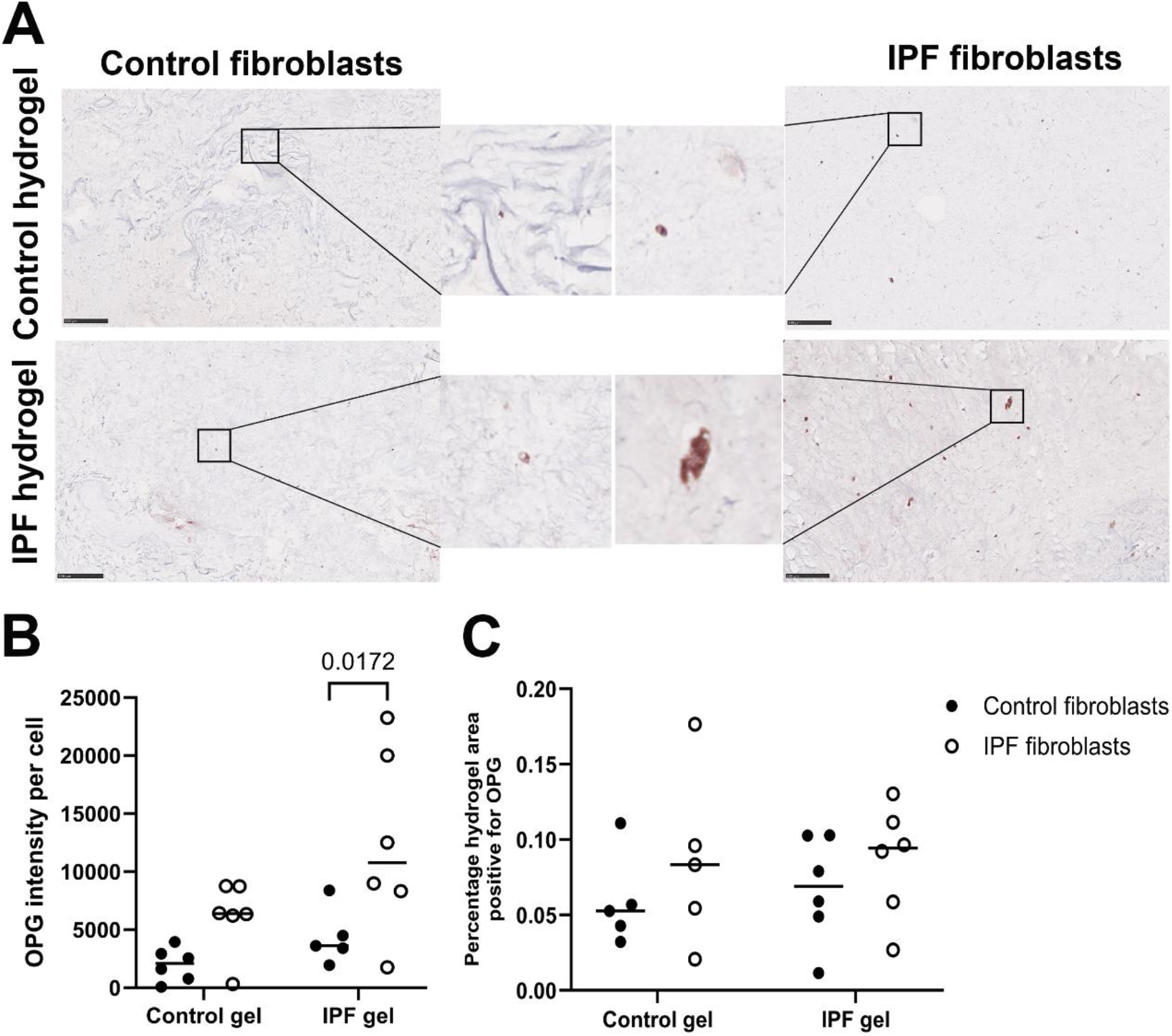
OPG staining in lung dECM-derived hydrogels encapsulated with fibroblasts. Control and fibrotic lung fibroblasts were cultured in control or IPF lung dECM-derived hydrogels for 7 days. The fixed hydrogel sections were stained to detect OPG deposition.(A) Representative images of staining for deposited OPG in control or IPF lung dECM-derived hydrogels encapsulated with control or IPF fibroblasts, nuclei are blue and OPG staining is red (scale bar = 250μm). (B) Quantification of total intensity of OPG per cell in fibroblasts encapsulated in control (n=5) and IPF (n=5 or 6) lung dECM hydrogel. (C) Percentage of pixels within the whole section image that met the criteria for the digital threshold for a strong positive signal for OPG in fibroblasts encapsulated in control (n=5) and IPF (n=6) lung dECM hydrogel. Data analysed using two-way ANOVA with Šidák-corrected post hoc test. P < 0.05 was considered significant, if no p values are mentioned in graphs values were not significant. OPG: osteoprotegerin, IPF: idiopathic pulmonary fibrosis, dECM: decellularized extracellular matrix.

### OPG correlated with stiffness, fiber length, fiber alignment and curvature when fibroblasts were in a fibrotic microenvironment

In our previous study we demonstrated that lung fibroblasts can modify matrix stiffness and structural properties of collagen fibres within a fibrotic microenvironment^11^, including the degree of high-density matrix, collagen fiber length, collagen fiber alignment, and curvature of collagen fibers. Combining these preceding data with our observations that fibrotic fibroblasts had enhanced capacity for release and deposition of OPG (Figures 1 and 2), we sought to examine whether the release and deposition of OPG were correlated with any of those biomechancial and structural properties. To this end, we evaluated correlations between secreted or deposited OPG with biomechanical, as well as structural properties. We observed correlations **only** in IPF hydrogels and not in control hydrogels (correlation data from control hydrogels are presented in supplemental Figures 1-4). Specifically, we found that when control fibroblasts were encapsulated in IPF hydrogels, they secreted less OPG when hydrogel stiffness increased (Figure 3A) and when collagen fiber length increased after 7 days of culture (Figure 3B). However, the amount of deposited OPG positively correlated with collagen fiber alignment (Figure 3C), while it trended towards a negative correlation with fiber curvature (Figure 3D). All parameters not specifically mentioned did not correlate with secreted or deposited OPG (supplement Figures 5-6). Strikingly, the correlations between secreted and deposited OPG and biomechanical and strucutral properties of the ECM exhibited reverse patterns when fibrotic fibroblasts were encapsulated in IPF hydrogels compared to control fibroblasts (Figure 3). Collectively, these data indicate that control and fibrotic lung fibroblasts reponded differently to a fibrotic microenvironment. These findings highlight that the microenvironment has a major influence upon lung fibroblast behavior and possibly this is enacted, in part, through an interaction between OPG and ECM fibers.

**Figure 3:**
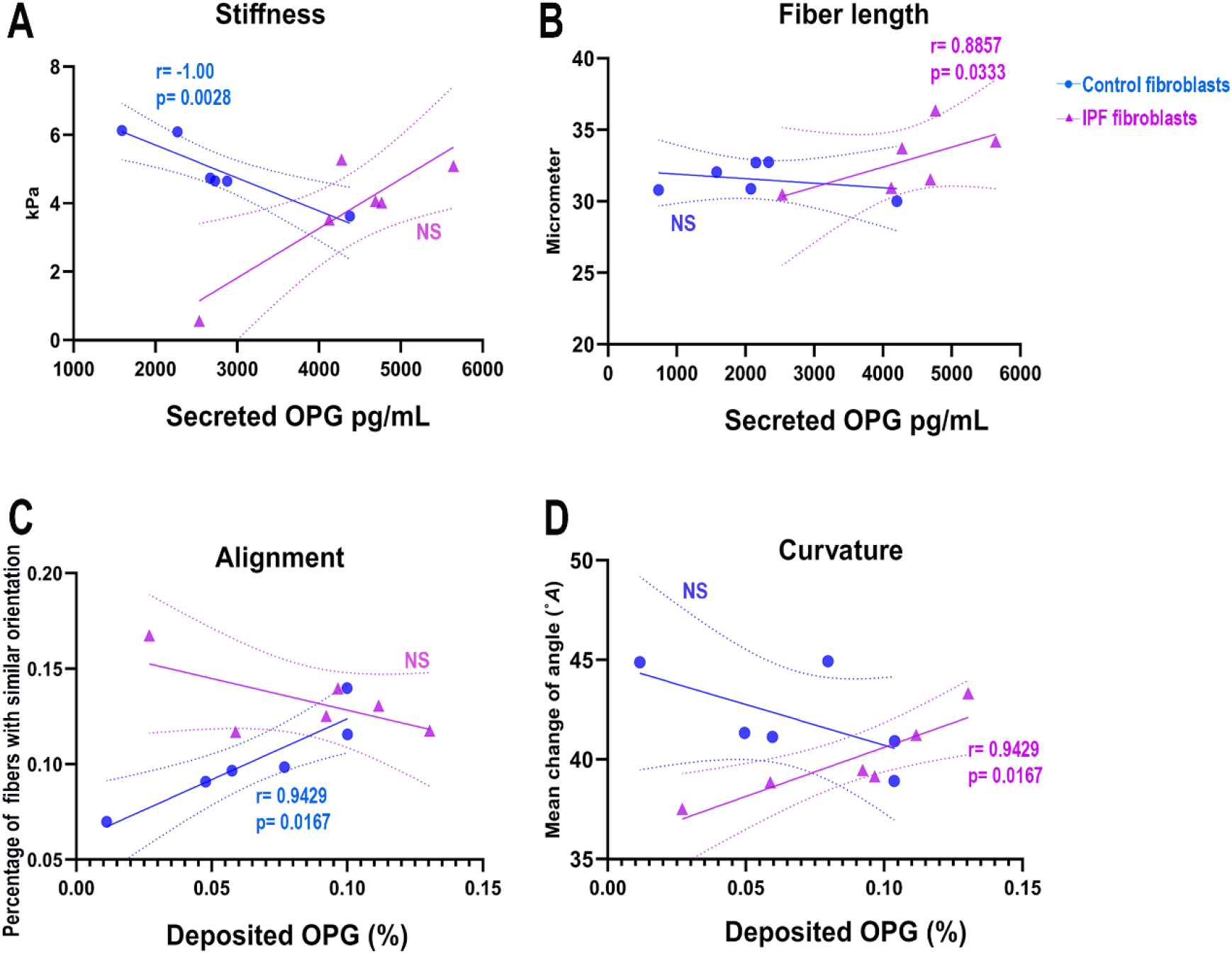
Correlations between OPG and biomechanical and structural ECM properties in IPF lung dECM hydrogels encapsulated with control and IPF fibroblasts. (A) Correlations between secreted OPG and stiffness in IPF lung dECM hydrogels encapsulated with control fibroblasts (n=6) or IPF fibroblasts (n=6). (B) Correlations between secreted OPG and fiber length in IPF lung dECM hydrogels encapsulated with control fibroblasts or (n=6) IPF fibroblasts (n=6). (C) Correlations between deposited OPG and alignment in IPF lung dECM hydrogels encapsulated with control fibroblasts (n=6) or IPF fibroblasts (n=6). (D) Correlations between deposited OPG and fiber curvature in IPF lung dECM hydrogels encapsulated with control fibroblasts (n=6) or IPF fibroblasts (n=6). Correlations were tested using a Spearman test, p< 0.05 was considered statistically significant, (0 < r ≤ 1) is positive correlation, and (−1 ≤ r < 0) is negative correlation. OPG: osteoprotegerin. OPG: osteoprotegerin, IPF: idiopathic pulmonary fibrosis, dECM: decellularized extracellular matrix.

### The association of OPG with the change in ECM stiffness is a response of lung fibroblasts to fibrotic microenvironment

Next we sought to investigate whether the presence of OPG in the matrix associated with the changes in ECM stiffness over 7 days in both control and fibrotic lung fibroblasts within control and IPF lung dECM hydrogels. Since this correlation was driven by these lung fibroblasts, we analyzed the relationship between cell-associated OPG levels and matrix stiffness changes. Notably, a correlation was observed only in IPF dCECM lung hydrogels again (Figure 4). In this context, OPG intensity per cell negatively correlated with stiffness change for control fibroblasts (Figure 4A), while for fibrotic fibroblasts this correlation between OPG intensity per cell and stiffness change was a significant positive association (Figure 4B). These findings suggest that control and fibrotic lung fibroblasts exhibit distinct responses to the fibrotic microenvironment, with OPG potentially playing a key role in associating with ECM stiffness dynamics in the fibrotic context.

**Figure 4:**
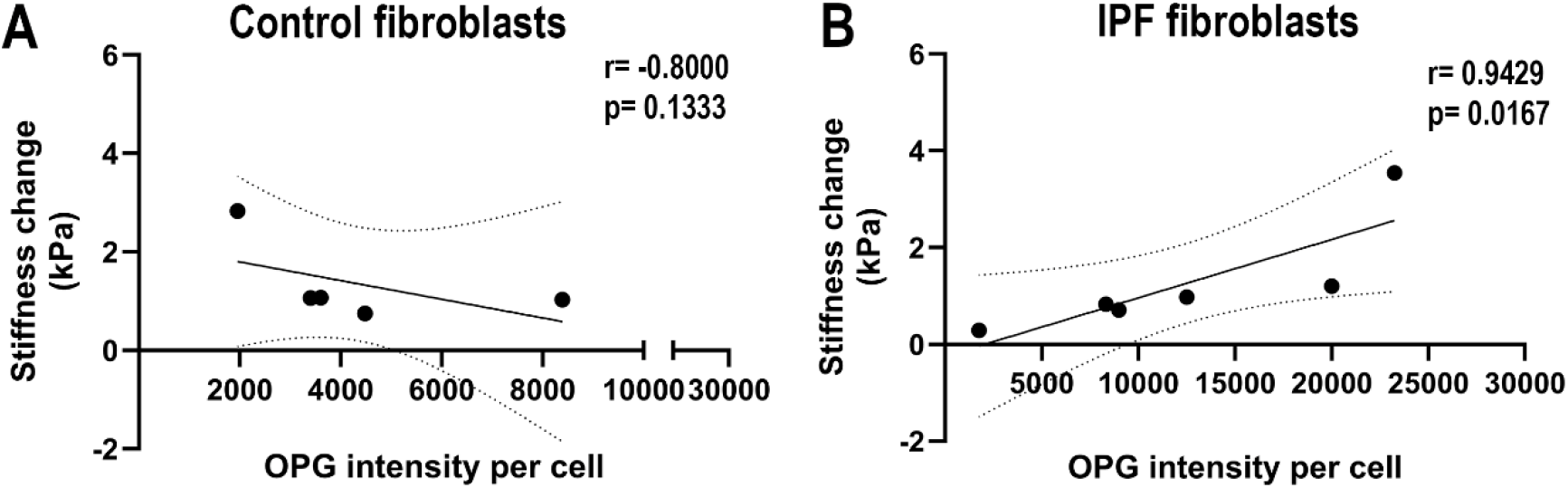
Correlations between OPG and changes in ECM stiffness in IPF lung dECM hydrogels encapsulated with control and IPF fibroblasts. Correlations between total intensity of OPG per cell and stiffness in IPF lung dECM hydrogels encapsulated with (A) control fibroblasts (n=5) or (B) IPF fibroblasts (n=6).Correlations were tested using a Spearman test, p< 0.05 was considered statistically signaficant, (0 < r ≤ 1) is positive correlation, and (−1 ≤ r < 0) is negative correlation. OPG: osteoprotegerin, IPF: idiopathic pulmonary fibrosis.

**Figure 5.**
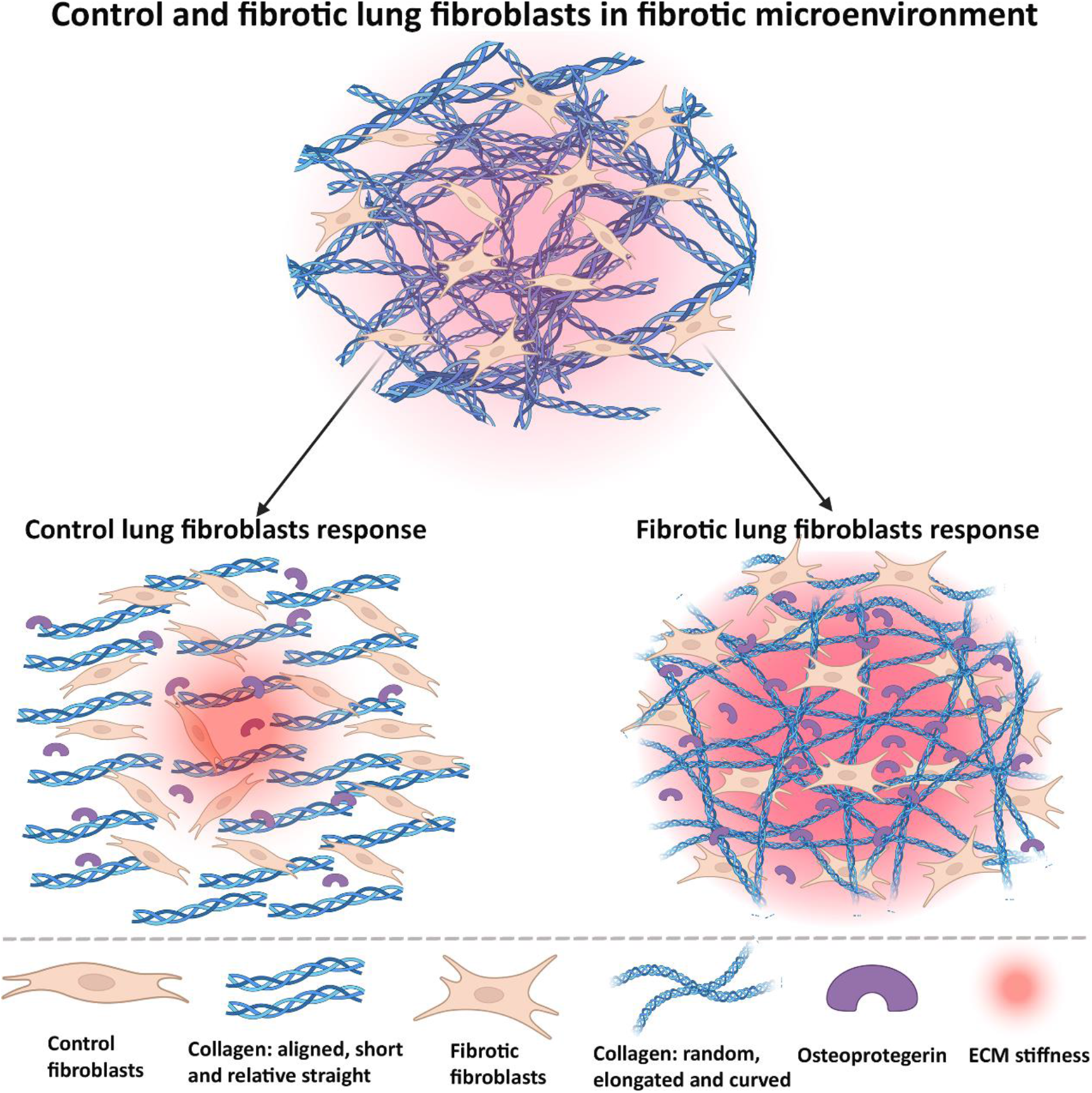
Summary schematic relationship of modulation of OPG, matrix stiffness and characteristics of collagen fibers in fibrotic microenvironments by control and fibrotic lung fibroblasts. In response to fibrotic stimulus, in control fibroblasts secreted OPG is negatively associated with matrix stiffness and deposited OPG positively with alignment of collagen fibers. The secreted OPG and deposited OPG from fibrotic fibroblasts were positively correlated with elongated and curvy collagen fibers. More OPG in control fibroblasts associates with low ECM stiffness; More OPG in fibrotic fibroblasts associates with high ECM stiffness and the changes in stiffness. OPG: osteoprotegerin; ECM: extracellular matrix.

## Discussion

In this study, we observed that a fibrotic microenvironment impacts the behavior of control and fibrotic fibroblasts in a different way. Within a fibrotic matrix microenvironment, for control fibroblasts a higher matrix stiffness is associated with lower presence of soluble OPG, suggesting retention of OPG in the matrix or lower production. This then coincided with the finding that when more OPG was retained in the fibrotic matrix, fibers became more aligned and less curved. Interestingly, an almost opposite pattern was found for fibrotic fibroblasts. For fibrotic fibroblasts in a fibrotic matrix context, more elongated collagen fibers were associated with higher presence of soluble OPG. It appeared that as more OPG was deposited into the matrix, the collagen fibers became curvier. Furthermore, we found that more OPG produced by control fibroblasts may lead to a decrease in matrix stiffness, whereas in fibrotic fibroblasts, OPG levels and ECM stiffness increased together in fibrotic microenvironment. This was corroborated by the finding that for control fibroblasts OPG intensity per cell correlated negatively with the change in stiffness experienced by the fibroblasts as these cells remodeled their environment, whereas with fibrotic fibroblasts this correlation between OPG intensity per cell and stiffness change maintained a positive correlation despite the cells remodeling their environment in a profibrotic manner. Altogether, these findings illustrate that the fibrotic microenvironment through mechanical stimuli drives control fibroblasts to dynamically and specifically respond by changing both OPG production and ECM structures and changes in ECM stiffness. This controlled dynamic reorganization of the fibrotic microenvironment is disrupted in fibrotic fibroblasts, potentially pointing to this mechanism being a driver of progression of fibrosis.

It is well-established that a fibrotic environment significantly influences cellular behavior. For instance, elevated fibrosis-associated gene expression was found in primary fibrotic fibroblasts cultured on decellularized fibrotic ECM derived from lung tissue of patients with IPF^23^. Fibroblasts cultured on collagen I gels develop large actin microfilament bundles and organize fibronectin into extracellular fibrils via transmitting stress to the collagen scaffold^24^. These findings indicate the importance of considering the influence of a pathological environment on fibroblast behavior in the context of fibrosis. In our study, we observed non-fibrotic and fibrotic fibroblasts produce OPG regardless of the environment. However, the balancing of the level of OPG and ECM remodeling orchestrated by fibroblasts was only evident within the fibrotic environment. This indicates that despite the fact that a fibrotic environment does not directly modulate the release of OPG from lung fibroblasts, the environment importantly influences release and deposition of OPG into the extracellular space, resulting in an association between OPG and ECM remodeling. OPG is a profibrotic factor that can be anchored by fibulin-1 into (fibrotic) lung matrix^25^. It can also directly bind to heparan sulfate glycosaminoglycans^26^, an ECM component regulating ECM fiber assembly^27^. It is plausible that OPG released from lung fibroblasts may be anchored by fibulin 1 into the fibrotic lung matrix. Subsequently, this matrix-associated OPG may be bound to a specific site of collagen (or other ECM components) in the matrix.

Control fibroblasts appear to exhibit a wound healing response in the fibrotic microenvironment. They regulate OPG deposition/secretion to modulate ECM stiffness and structure, aiming to restore the fibrotic microenvironment toward a normal nonfibrotic state, while fibrotic fibroblasts may have adapted to their fibrotic environment and have lost their ability to return to homeostasis through ECM remodeling. This distinction may be due to the origin of lung fibroblasts. Robin^28^ and Fereshteh et al^29^ summarized the origin and function of fibroblasts in disease conditions, different phenotypes of fibroblasts exhibit various distinct functions in a diseased microenvironment. This may support the data in our study, which is that control and fibrotic lung fibroblasts exhibit an opposite behavior in response to fibrotic stimuli by orchestrating alterations of ECM stiffness and the amount of OPG. However, the mechanism driving fibroblasts’ distinct behaviors remains underinvestigated. Our data suggest that this may be due to alternate sensing of mechanical cues by control and fibrotic fibroblasts. In support of this, control primary lung fibroblasts can actively increase Piezo1 protein levels, a mechanosensitive protein, when mechanically stretched^30^. Notably, activated Piezo1 has been demonstrated to upregulate OPG protein expression in control osteocytes^31^ and a lack of Piezo1 could suppress OPG gene expression in osteoblasts^32^. Further investigations can explore whether mechanosensitive factors have an impact on the association between OPG and ECM remodeling facilitated by fibroblasts in fibrotic microenvironment.

Our findings highlight novel patterns in the association of OPG with matric stiffness and collagen organization in both control and fibrotic fibroblasts within the fibrotic microenvironment. However, the role of OPG in lung fibrosis remains unclear. It is still unknown whether OPG directly influences ECM remodeling and stiffness, or it indirectly affects the alterations in ECM by acting as a modulator of other underlying processes. Future studies should focus on the mechanisms by which OPG interacts with changes in ECM in lung fibrosis.

## Conclusions

Overall, this study indicates that control and fibrotic lung fibroblasts respond differently to a fibrotic microenvironment. These two types of lung fibroblasts exhibit opposite behaviors in response to this fibrotic stimulus. This difference is seen in the ECM remodeling process in terms of the quantity of secreted/deposited OPG associated with mechanical, as well as structural ECM characteristics (Figure 5). Further investigations should focus on what drives control and fibrotic lung fibroblasts to exhibit these different behaviors, as well as the mechanisms underlying how fibroblasts modulate OPG together with ECM stiffness, ECM structural characteristics and vice versa in fibrotic lung environments. Such investigations may pave a novel way for better understanding of the development and progression of lung fibrosis.

## Notes

### Competing Interest Statement

The authors have declared no competing interest.

