## Supplemental Figure 1, Supplemental Figure2, Supplemental Figure3, Supplemental Figure 4, Supplemental Figure5, Supplemental Figure6 for "Contrasting responses of control and fibrotic lung fibroblasts to fibrotic stimuli: the role of osteoprotegerin in extracellular matrix remodeling"

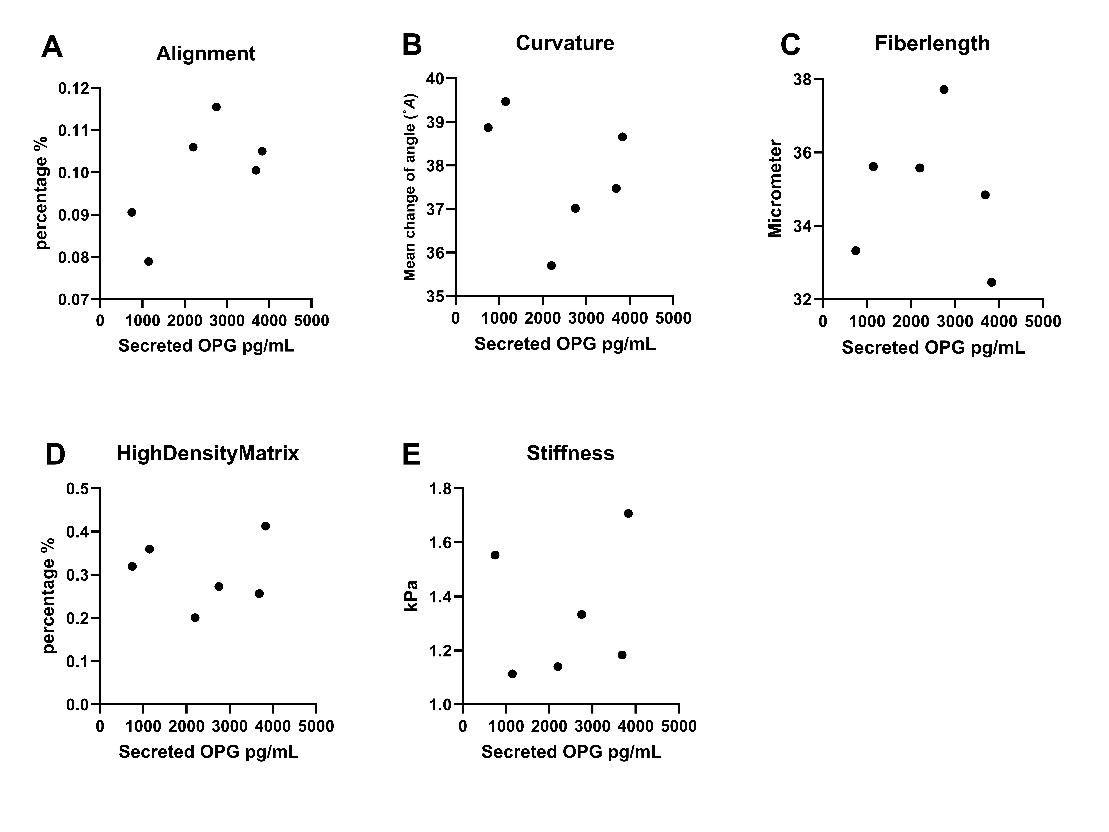

**Figure 1: Correlations between secreted OPG and biomechanical and structural ECM properties in control hydrogels seeded with control fibroblasts (n=6).** Correlations between secreted OPG with (A) fiber alignment (r=0.4857, p=NS), (B) fiber curvature (r=-0.3714, p=NS), (C) fiber length (r=-0.2571, p=NS), (D) high density matrix (r=0.0857, p=NS) and (E) matrix stiffness (r=0.3714, p=NS). Correlations were calculated using a Spearman test.

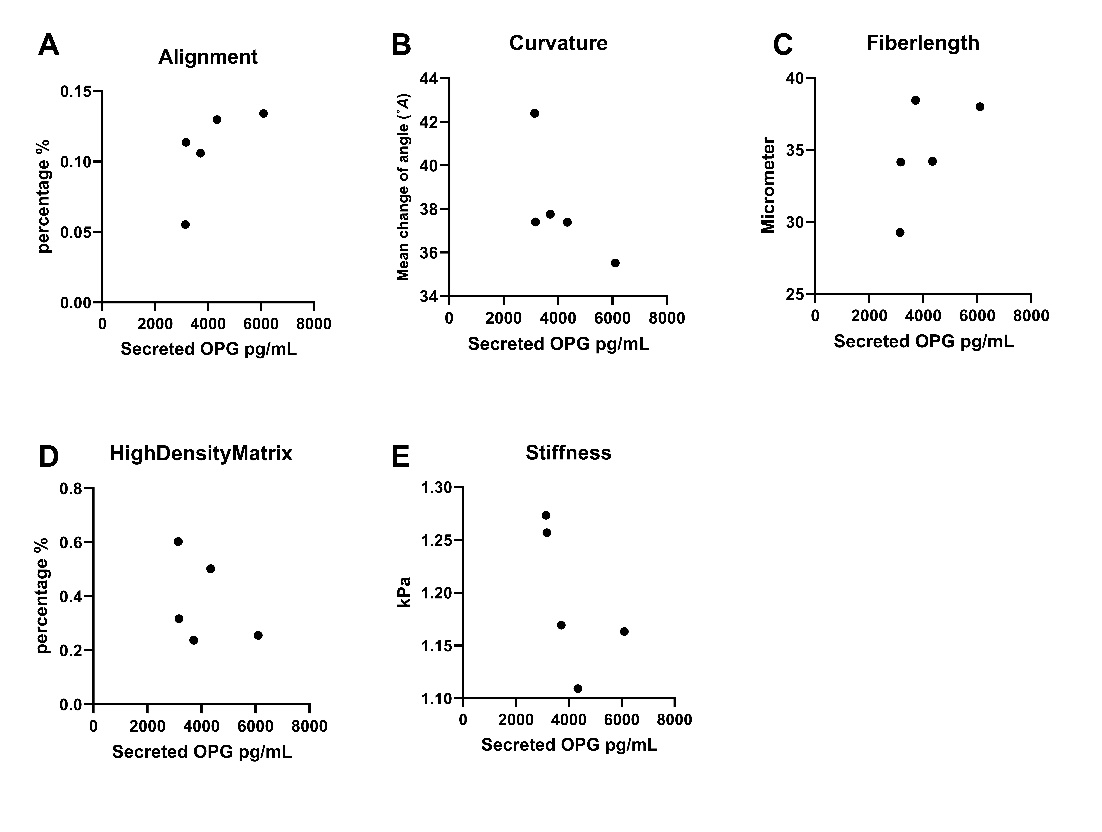

**Figure 2: Correlations between secreted OPG and biomechanical and structural ECM properties in control hydrogels seeded with IPF fibroblasts (n=5).** Correlations between secreted OPG and stiffness in IPF hydrogels seeded with control fibroblasts Correlations between secreted OPG with (A) fiber alignment(r=0.900, p=0.0833), (B) fiber curvature (r=-0.900, p=0.0833), (C) fiber length (r=0.700, p=NS), (D) high density matrix (r=-0.500, p=NS) and (E) matrix stiffness (r=-0.900, p=0.083). Correlations were calculated using a Spearman test.

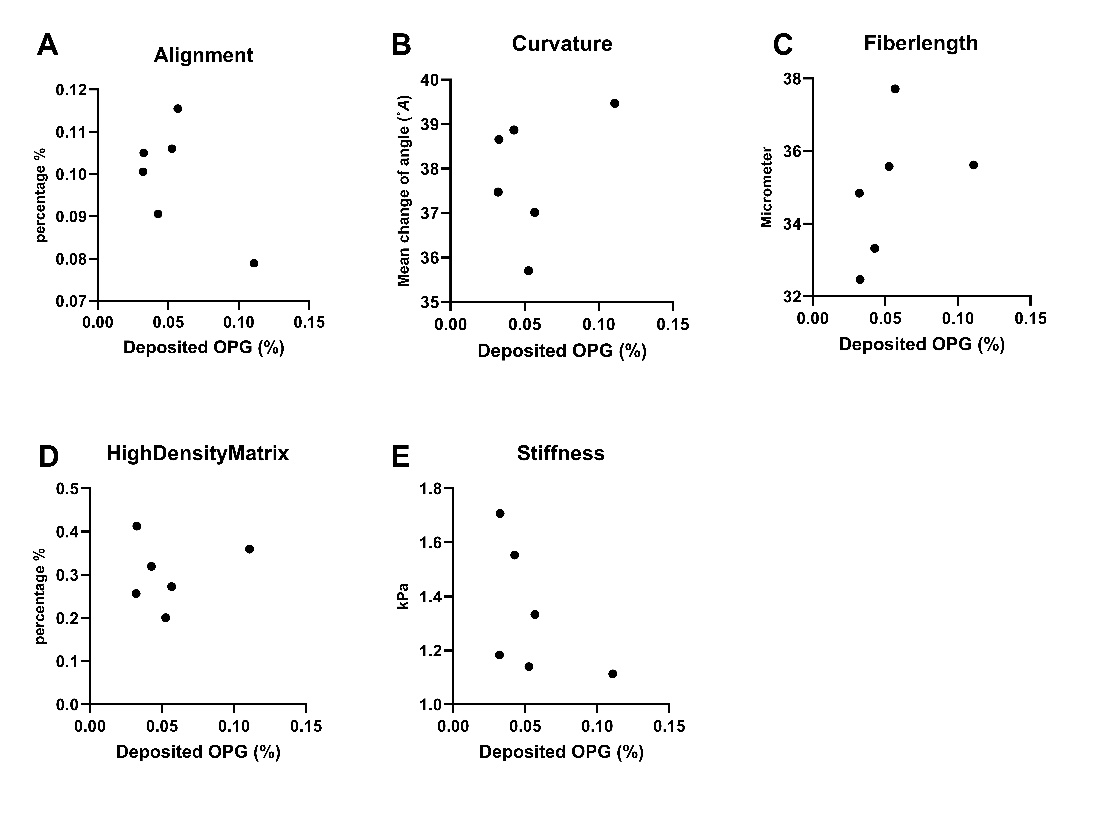

**Figure 3: Correlations between deposited OPG and biomechanical and structural ECM properties in control hydrogels seeded with control fibroblasts.** Correlations between secreted OPG with (A) fiber alignment (r=-0.0286, p=NS), (B) fiber curvature (r=0.1429, p=NS), (C) fiber length (r=0.7714, p=NS), (D) high density matrix (r=0.0857, p=NS) and (E) matrix stiffness (r=-0.5429, p=NS). Correlations were calculated using a Spearman test.

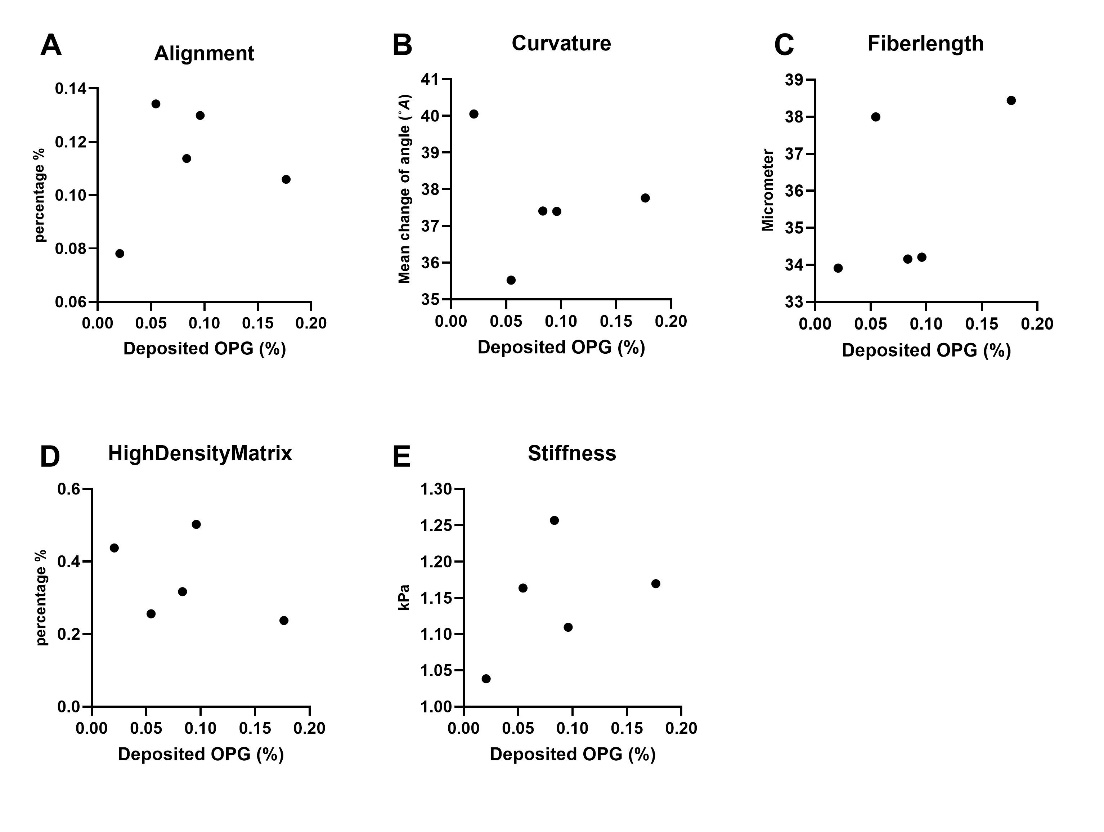

**Figure 4: Correlations between deposited OPG and biomechanical and structural ECM properties in control hydrogels seeded with IPF fibroblasts.** Correlations between secreted OPG with (A) fiber alignment (r=0.100, p=NS), (B) fiber curvature (r=-0.100, p=NS), (C) fiber length (r=0.700, p=NS), (D) high density matrix (r=-0.300, p=NS) and (E) matrix stiffness (r=0.500, p=NS). Correlations were calculated using a Spearman test.

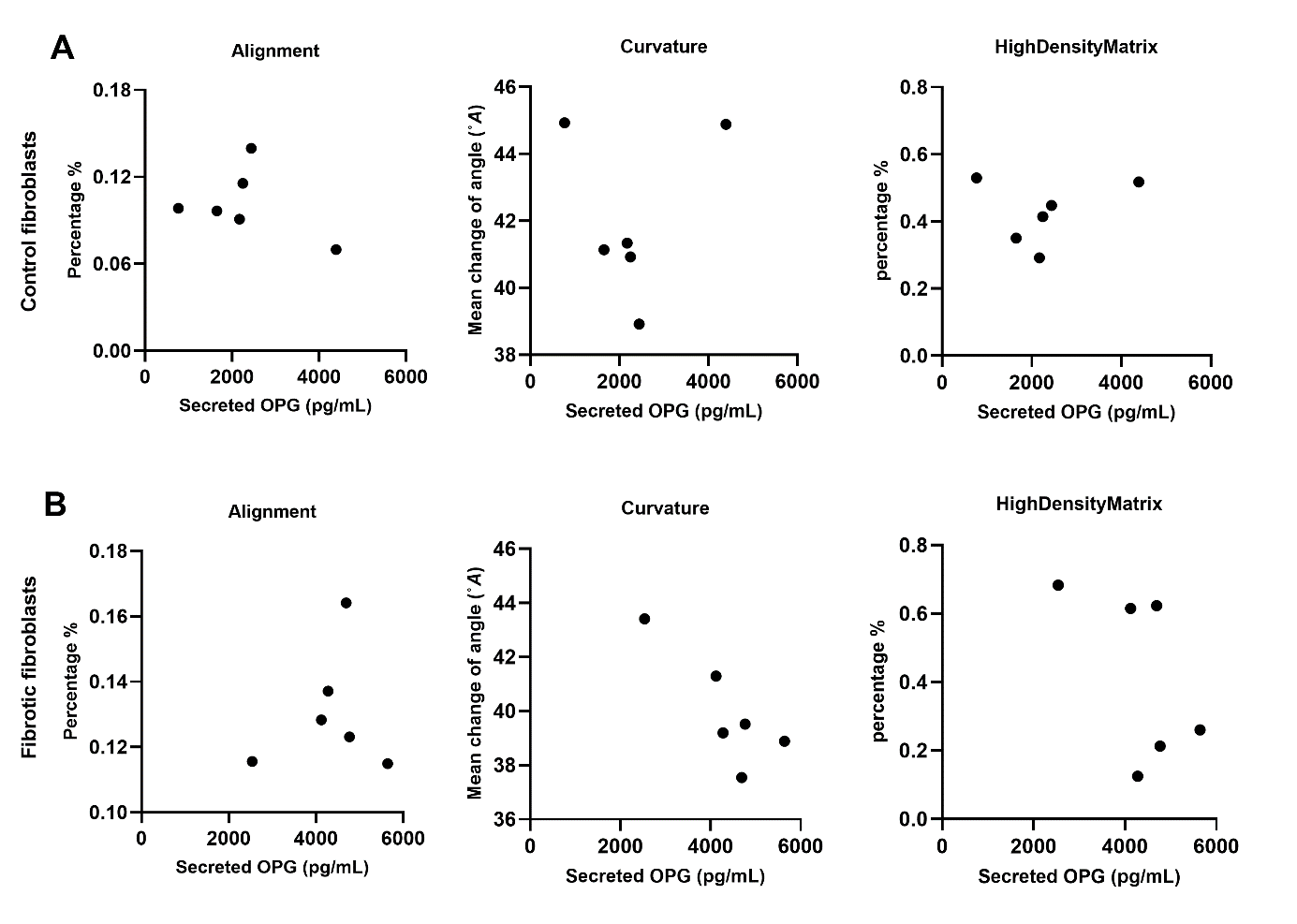

**Figure 5: Correlations between secreted OPG and biomechanical and structural ECM properties in IPF hydrogels seeded with control or fibrotic fibroblasts.** (A) Correlations between secreted OPG with fiber alignment (r=-0.08571, p=NS), fiber curvature (r=--0.3714, p=NS), high density matrix (r=0.08571, p=NS) for control fibrobalsts. (B) Correlations between secreted OPG with fiber alignment (r=-0.2000, p=NS), fiber curvature (r=-0.7143, p=NS), high density matrix (r=-0.4857, p=NS) for fibrotic fibrobalsts. Correlations were calculated using a Spearman test.

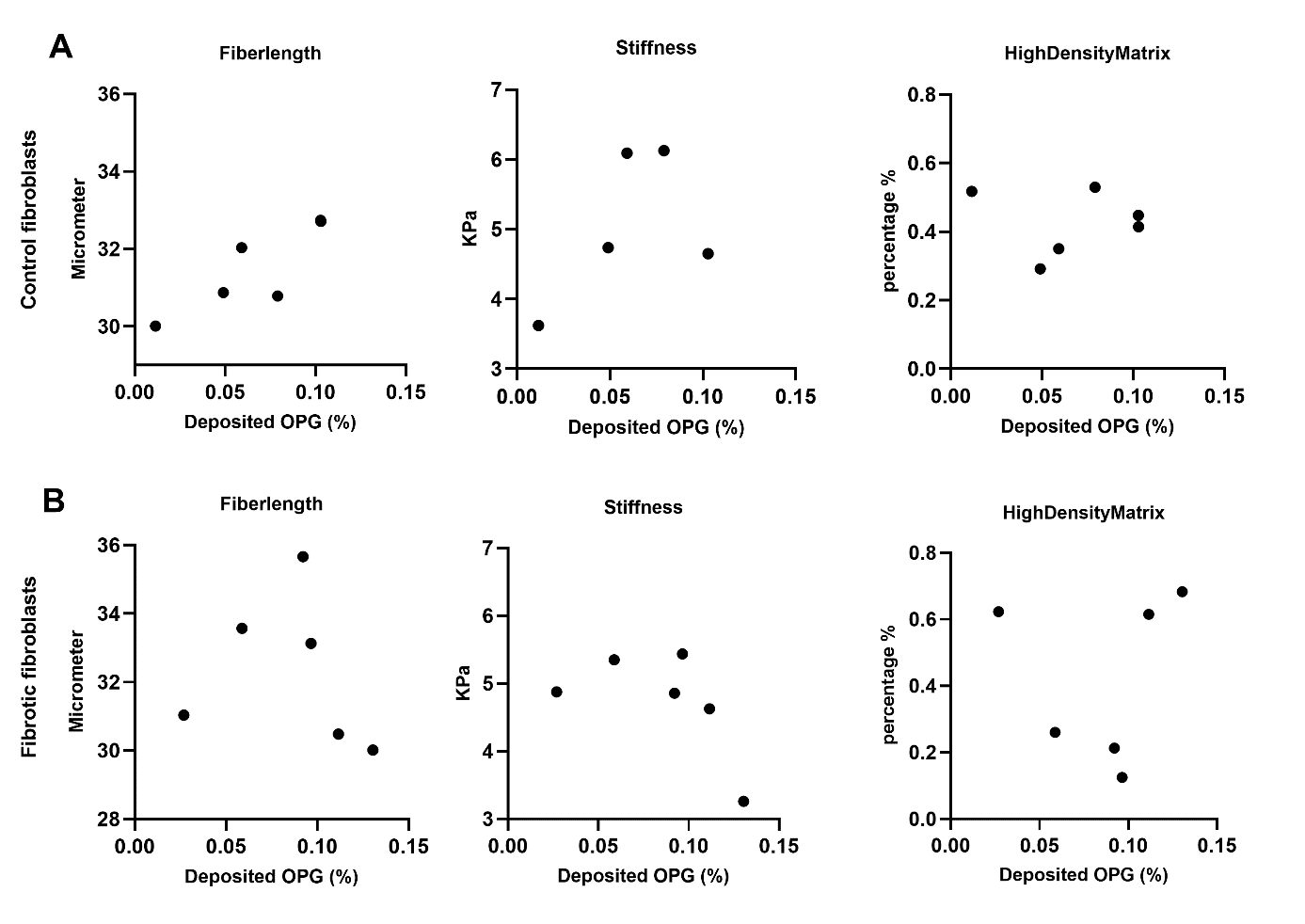

**Figure 6: Correlations between deposited OPG and biomechanical and structural ECM properties in IPF hydrogels seeded with control or fibrotic fibroblasts.** (A) Correlations between secreted OPG with fiber lengh (r=0.7714, p=NS), stiffness (r=0.1429, p=NS) and high density matrix (r=0.08571, p=NS) for control fibrobalsts. (B) Correlations between secreted OPG with fiber length (r=-0.6000, p=NS), stiffness (r=-0.600, p=NS) and high density matrix (r=0.200, p=NS) for fibrotic fibrobalsts. Correlations were calculated using a Spearman test.
